# Tissue mimetic hyaluronan bioink containing collagen fibers with controlled orientation modulating cell morphology and alignment

**DOI:** 10.1101/2020.02.26.966564

**Authors:** Andrea Schwab, Christophe Helary, Geoff Richards, Mauro Alini, David Eglin, Matteo D’Este

## Abstract

Biofabrication is providing scientists and clinicians the ability to produce engineered tissues with desired shapes, and gradients of composition and biological cues. Typical resolutions achieved with extrusion-based bioprinting are at the macroscopic level. However, for capturing the fibrillar nature of the extracellular matrix (ECM), it is necessary to arrange ECM components at smaller scales, down to the micron and the molecular level.

In this study, we introduce a bioink containing hyaluronan (HA) as tyramine derivative (THA) and collagen type 1 (Col 1). Similarly to other connective tissues, in this bioink Col is present in fibrillar form and HA as viscoelastic space filler. THA was enzymatically crosslinked under mild conditions allowing simultaneous Col fibrillogenesis, thus achieving a homogeneous distribution of Col fibrils within the viscoelastic HA-based matrix. THA-Col composite displayed synergistic properties in terms of storage modulus and shear-thinning, translating into good printability.

Shear-induced alignment of the Col fibrils along the printing direction was achieved and quantified via immunofluorescence and second harmonic generation. Cell-free and cell-laden constructs were printed and characterized, analyzing the influence of the controlled microscopic anisotropy on human bone marrow derived mesenchymal stromal cells (hMSC) migration.

THA-Col showed cell instructive properties modulating hMSC adhesion, morphology and sprouting from spheroids stimulated by the presence and the orientation of Col fibers. Actin filament staining showed that hMSCs embedded into aligned constructs displayed increased cytoskeleton alignment along the fibril direction. Based on gene expression of cartilage/bone markers and ECM production, hMSCs embedded into the bioink displayed chondrogenic differentiation comparable to standard pellet culture by means of proteoglycan production (Safranin O staining and proteoglycan quantification).

The possibility of printing matrix components with control over microscopic alignment brings biofabrication one step closer to capturing the complexity of native tissues.

## 1 Introduction

Biofabrication aims at engineering constructs recapitulating the complexity of mammal tissues concerning cell types, chemical and biological gradients, and multiscale architecture. In extrusion-based 3D printing (3DP), the resolution is determined by the size of the nozzle, printing speed and offset, distance between nozzle and printing surface, with resolution ranging from the mm down to the micron range [1]. The physico-chemical properties of the biomaterial and bioink directly influence printing outcome and shape fidelity. The ink is most often a viscoelastic shear-thinning hydrogel (precursor) with good elasticity recovery after high shear and rapid gelation after extrusion [2-4]. In 3DP, high resolution must be compromised for cell viability, with typical resolutions in the hundreds of microns range [5]. Thus, for standard extrusion-based techniques, which are the most versatile and widespread, only the macroscopic architecture can be deliberately designed.

By combining different biomaterial ink compositions, gradients of material composition and cellular distribution can be produced using 3D printing techniques [6-8]. Despite impressive advances, the field still lacks reliable methods to mimic tissue architectures not only by replicating the ECM composition, but also addressing the (macro)molecular organization within the biomaterial ink at (sub-) micron scale lengths, *e.g*. on fibrillar levels [9]. However, it is well-known how microarchitectural features provide specific and unique properties to natural tissues [10].

Col is the most abundant protein in the ECM, where it is found with a hierarchical fibrillar structure ranging from the triple helix at the molecular level, up to the fibrils and fibers on the microscopic level. Col structure, orientation and spatial arrangement are fundamental towards mechanical stability and anisotropic properties of tissues [11]. The network architecture is also crucial to transmit forces to cells via cell-matrix interaction and contributes to the matrix biochemical environment [12, 13].

To bring the 3DP technology one step closer to native tissue architectures, it is necessary to capture micro- and nanostructures within macroscopically complex scaffold geometries [14]. Microarchitectural properties can be introduced with polymer self-assembling, a process where macromolecules arrange into stable non-covalent structures. Col 1 [15], fibrin [12], cellulose [16] and silk fibroin [17] are the most prominent materials self-assembling into fibrous structures.

Most studies printing Col 1 biomaterial inks employ soluble (acidic) and/or non-fibrillar Col 1 inks or blended neutralized Col 1 (Col 1 fractions). Printing an acidic or neutral Col 1 dispersion (20-40 mg/ml) of swollen Col fibrils with a high viscosity (>10^3^ P·s at shear rate of 0.1 1/s) requires subsequent lyophilization and chemical crosslinking for scaffold stabilization [18]. Neutralized non fibrillar Col 1 (8 mg/ml) was held at low temperature before extrusion and gelation was induced by printing at 37°C [19]. In another study, neutralized Col 1 fractions were printed at high concentrations (20-40 mg/ml) and lower temperature (4-15°C) during extrusion printing and gelled within pre warmed media [20].

However, only few studies investigated the formation and presence of Col 1 fibers within the printed hydrogel construct and the influence on cell migration [21-23]. Moreover, cell embedding is limited in the above-mentioned approaches either due to the acidic conditions or the post processing. Another drawback of printing neutralized Col 1 is the phase separation of the self-assembled fibrils from the liquid when extruded [23].

Hyaluronic acid or hyaluronan (HA) is another abundant ECM component osmotically capable of holding large amounts of water, and thus functioning as a space filler for fibrillar matrix components, thus providing compressive strength through fluid retention [24]. To our knowledge, methods to control the orientation of Col 1 fibers within a viscoelastic HA-based matrix for bio-fabrication has not been addressed before.

The aim of this study was the development of a THA-Col 1 composite biomaterial ink and the workflow to 3D print it, fabricating constructs with homogeneous composition and controlled Col fibril orientation at the microscopic level. In addition, the impact of the microscopic anisotropy on the behavior of the embedded hMSC was analyzed (Fig. 1). To this aim, the tyramine derivative of HA (THA) was employed to form a continuous matrix where solubilized Col 1 was homogeneously dispersed; THA gelation and Col fibrillogenesis occurred without mutual interference. THA was additionally crosslinked with visible light for final shape fixation, thus achieving a uniform THA matrix containing Col fibrils, which were aligned via the shear forces experienced during the extrusion process. Fibril alignment and cytoskeleton orientation were quantified and compared to casted, isotropic constructs. hMSC spheroids were embedded in isotropic constructs with different compositions of THA and Col to compare their sprouting and chondrogenic differentiation.

**Figure 1:**
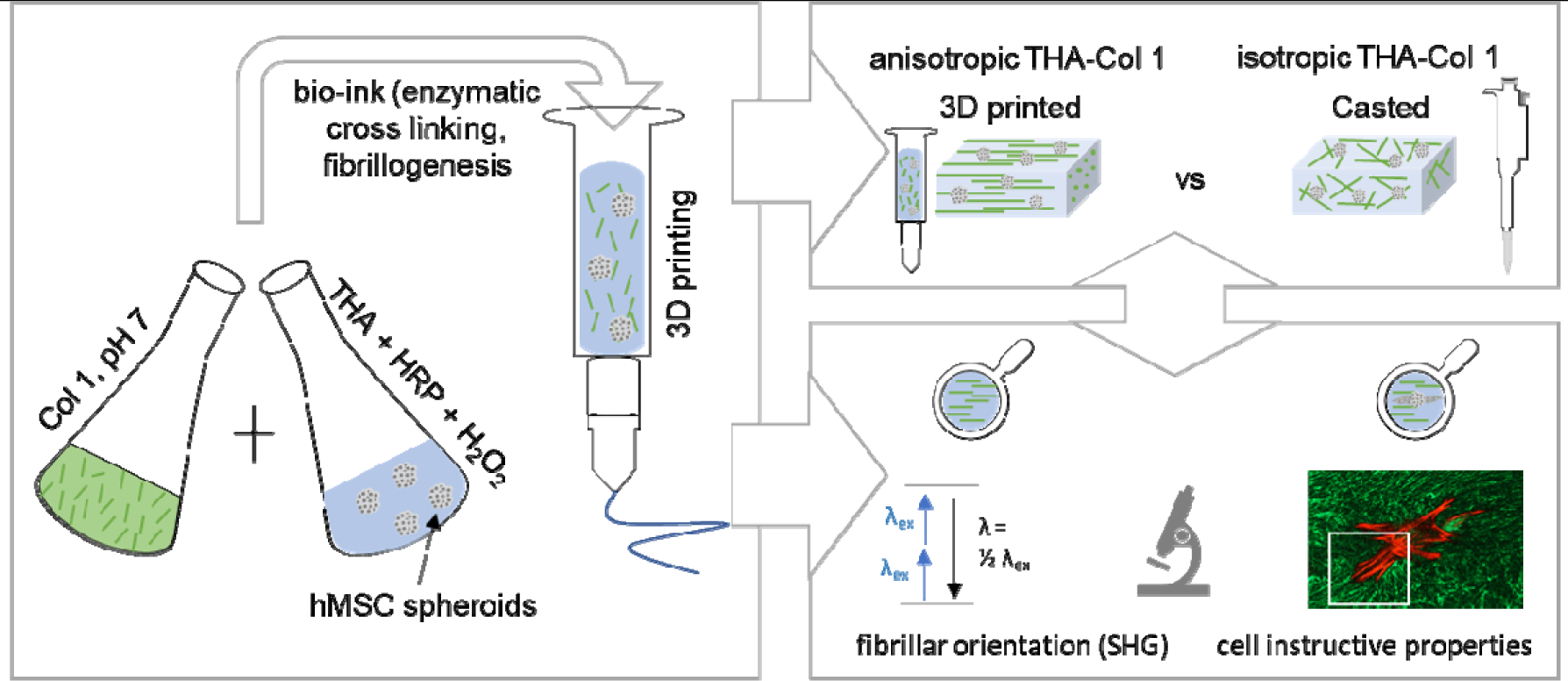
3D bioprinting as a tool to produce microscopic anisotropic scaffolds. Biomaterial and bioink were prepared by mixing neutralized Col 1 (5 mg/ml) isolated from rat tails with tyramine modified HA (THA, 25 mg/ml). Neutralized Col 1 was mixed with THA for enzymatic crosslinking either cell free or containing hMSC cell spheroids. The Col 1 microstructure was investigated after 3D printing and compared to casted, isotropic samples with different microscopic techniques (Second Harmonic Generation SHG imaging and confocal microscopy). Cell instructive properties were analyzed after in vitro culture on cell migration and attachment.

## 2 Materials and Methods

### 2.1 Tyramine modified hyaluronic acid (THA) synthesis

THA was synthesized as previously described [25]. Briefly, HA (280-290 kDa, 5 mM carboxylic groups, Contipro Biotech S.R.O) was functionalized via 4-(4,6-dimethoxy-1,3,55-triazin-2-yl)-4-mehtylmorpholinium chloride (DMTMM, TDI) amidation with tyramine by mixing with a stoichiometric ratio of 1:1:1. Functionalization was performed for 24h at 37°C. THA was precipitated by dropwise adding Ethanol (96% v/v), isolated with Gooch filter No. 2 and dried. The degree of substitution was 6.6% as determined by absorbance reading at 275 nm (Multiskan™ GO Microplate Spectrophometer, Thermo Fisher Scientific).

### 2.2 THA-Col 1 hydrogel preparation

THA (25 mg/ml) was reconstituted with minimum Essential Medium alpha (α-MEM, Gibco) supplemented with 10% v/v bovine fetal serum containing peroxidase from horseradish (HRP, Sigma Aldrich) at the specified concentration at 4°C under agitation. THA for turbidity measurement was reconstituted with PBS. Enzymatic crosslinking was initiated by adding hydrogen peroxide (H_2_O_2_, Sigma Aldrich) in different concentrations and incubated for 30 min at 37°C. THA-Col 1 composite was prepared by neutralizing Col 1 (5 mg/ml, Collagen I rat tail in 0.2 N acetic acid, Corning) with NaOH and adding to THA before enzymatic crosslinking was initiated. Mixing ratios by volumes of THA (25 mg/ml) with Col 1 (5 mg/ml) are given in brackets for all experimental set ups described in the following paragraphs. For cell encapsulation, hMSC spheroids were resuspended with THA before adding H_2_O_2_. Additional light crosslinking using 0.2 mg/ml of Eosin Y as photoinitiator (Sigma Aldrich) was done for all printing experiments to stabilize the printed structures and the cell migration and cell differentiation studies. Non-aligned constructs were produced by pipetting the hydrogel precursor with a positive displacement pipette (CP1000, inner diameter ID 1.0 mm, Gilson) into custom made silicon molds (6 mm in diameter). Enzymatic crosslinking was performed for all samples for 30 min at 37°C.

### 2.3 3D bioprinting of THA-Col

An extrusion based bioprinter (3D Discovery™, RegenHU) was used to prepare anisotropic hydrogels (0.25” cylindrical needles, 15G: ID 1.36 mm, 0.2 bar, writing speed 8 mm/s; 18G: ID 0.84 mm, 1.6 bar, writing speed 8 mm/s, Nordson EFD) with subsequent light crosslinking (515 nm LED, speed 4 mm/s). Biomaterial and bioink were transferred into 3CC barred (ID 2.3 mm, Nordson) for enzymatic crosslinking (30 min, 37°C), printed with the above-mentioned parameters and transferred into culture media (cell embedded hydrogel) or PBS (cell free).

A complex microstructure to mimic microarchitecture of Col fibers in articular cartilage was accomplished with CAD Software (RegenHU). Fibrillar structure in the superficial zone (SZ at the surface) was realized by printing 3 layers of parallel lines, whereas the Middle (MZ) and deep zone (DZ) were designed as circular structures to mimic the arch like geometry.

### 2.4 Rheological characterization

Viscoelastic properties of the biomaterial inks were analyzed using Anton-Paar MCR-302 rheometer with 1° cone-plate geometry and gap distance of 0.049 mm at 20°C. Silicone oil (Sigma Aldrich) was applied to the external border to prevent drying during the measurement. Viscosity was measured with shear rate ranging from 0.01 1/s to 100 1/s (n=2/group) to evaluate shear thinning behavior of the enzymatically crosslinked THA (0.3 U/ml HRP, 0.52 mM H_2_O_2_), THA-Col 1 (1:1 v/v) and Col 1 in a rotational experiment. Oscillatory tests (amplitude sweep: frequency 1 Hz, amplitude 0.01 - 100% strain; frequency sweep: amplitude 1% strain, frequency 0.01 - 100 Hz, n=2/group) were performed with parallel plate measuring system at 20°C to characterize the elastic modulus of THA and THA-Col 1 at varying polymer concentrations (0.3 U/ml HRP, 0.65 mM H_2_O_2_).

### 2.5 Turbidity measurement

Fibrillation of Col 1 was evaluated by absorbance reading at 313 nm at 37°C at 5, 30 and 60 min after induction of neutralization of Col 1 and enzymatic gelation (n=2-5 samples/group) of freshly prepared materials with a Multiskan™ GO Microplate Spectrometer (Thermo Scientific). The following solutions were analyzed: neutralized Col 1, acidic Col 1, THA (0.075 U/ml HRP, 0.26 mM H_2_O_2_) and THA-Col 1 (1:1 and 1:2 v/v) composite undergoing enzymatic cross-linking during measurement of Col 1 fibrillogenesis. Temperature was held at 37°C to induce enzymatic crosslinking of THA and stabilize Col 1 fibrillation.

### 2.6 Second harmonic generation (SHG) imaging

To visualize Col 1 fibers (0.3 U/ml HRP, 0.52 mM H_2_O_2_) (1:1 v/v) SHG images were acquired with a MaiTai multi-photon laser equipped confocal microscope (Leica SP8). SHG signal was collected with HyD detector between 437-453 nm (λ_ex_ 880 nm, output power ap 1.7 W). Additionally, transmitted light and autofluorescence signal (510 - 600 nm) was acquired with varying emission wavelength for SHG detector (λ_em_ 420-436 nm and λ_em_ 454-470 nm) to check for the specificity of SHG signal at 437-453 nm. For deep imaging a long-distance objective (25x water-immersion objective) was used to acquire z-stacks with optimal settings for each sample. Image series acquired with Leica Application Suite X software (LAS X, Leica) were processed with ImageJ (National Institute of health, NIH).

### 2.7 Cell culture

#### 2.7.1 Cell isolation

hMSCs were isolated from bone marrow aspirate with full ethical approval (Ethics committee of University of Freiburg Medical Centre - EK-Freiburg: 135/14) as described elsewhere [26]. hMSCs were sub-cultured with α-MEM (Gibco) supplemented with 10% v/v Sera Plus bovine serum (PAN Biotech), 100 U/ml Penicillin, 100 ug/ml Streptomycin (Gibco) and 5 ng/ml basic Fibroblast Growth Factor (FGFb, Fitzgerald Industries International) in a humidified atmosphere of 5% CO_2_ with media change every second day.

#### 2.7.2 Cell migration and viability study

Cell spheroids were prepared from hMSC (passage 3-4) in ultra-low attachment petri dish (Corning) with α-MEM media supplemented with 10% bovine serum and 100 U/ml Penicillin, 100 ug/ml Streptomycin before embedded in hydrogels. Cell spheroid suspension was added for (enzymatic) crosslinking (0.3 U/ml HRP, 0.39 mM H_2_O_2_) into THA-Col 1(1:1 v/v) or THA or Col 1 at final cell density of 3 Mio/ml and cultured for 8 days. For THA-Col 1 and THA, an additional light crosslinking of printed and casted samples was done at 515 nm.

To evaluate cell migration, samples at day 0, 3, and 8 were stained with phalloidin as described in 2.8.1 and fluorescent images were taken for subsequent quantification of migration length and area (n=3-5 spheroids/timepoint) described in Chapter 2.9.

Live and Dead assay was performed during the migration experiment of hMSC spheroids embedded in THA-Col 1 (0.3 U/ml HRP, 0.39 mM H_2_O_2_, 1:1 v/v) of 3D printed and casted samples at day 1 and day 6. H_2_O_2_ concentration was selected in a range known not to be cell toxic [27]. After washing with PBS, samples were incubated with 2 µM Calcein AM (Sigma Aldrich) and 1 µM Ethidium homodimer-1 (Sigma Aldrich) for 30 min at 37°C, washed with PBS and imaged using confocal microscope (LSM800, Carl Zeiss). Dead cells were stained with Ethidium homodimer-1 in red (λ_ex_ 561 nm), cytoplasm of living cells was stained with Calcein-AM in green (λ_ex_ 488 nm).

#### 2.7.3 Chondrogenic differentiation

For chondrogenic differentiation, hMSC (passage 2) were seeded in 6 well microwell plate (AggreWellTM 400 plate) at 1.67 Mio/well for 3 days to from cell spheroids. Chondrogenic media was composed of high glucose Dulbecco’s Modified Eagle Medium (DMEM HG, Gibco) supplemented with non-essential amino acids (1% v/v, Gibco), ascorbic acid 2 phosphate (50 ug/ml, Sigma Aldrich), dexamethasone (100 nM, Sigma Aldrich), ITS+ premix (1% v/v, Corning), TGF β1 (10 ng/ml, Fitzgerald) and antibiotics (10 U/ml Penicillin, 10 ug/ml Streptomycin, Gibco). Cell spheroids were embedded in THA-Col 1 (0.5 U/ml HRP, 0.65 mM H_2_O_2_, 5% w/v Col 1, 1:1 v/v) at final cell concentration of 5 Mio/ml hydrogel volume (400,000 cells/scaffold) before enzymatic gelation, additionally light cross linked, and cultured for 21 days with chondrogenic media. Media was changed 3 times a week. For positive control, standard hMSC pellets were prepared by seeding 250,000 hMSCs into each of a 96 V-Bottom well plate (Ratiolab) and cultured under same conditions.

### 2.8 Histological processing and staining

#### 2.8.1 Actin filament-Col 1 immunofluorescence staining

For actin filament staining in combination with Col 1 immuno fluorescent staining, printed and casted THA-Col 1 (1:1 v/v) were fixed with 4% neutral buffered formalin (Formafix AG) at room temperature and stored in PBS at 4°C upon staining.

Cytoskeletal organization of hMSC sprouting from cell spheroids embedded in enzymatically cross linked THA-Col 1 samples (0.3 U/ml HRP, 0.39 mM H_2_O_2_) were analyzed on day 0 and after 6 days culture with Phalloidin staining to visualize actin filaments. Hydrogels were permeabilized with 0.5% v/v Triton X-100 (Sigma Aldrich) for 10 min at room temperature, blocked with 10% bovine serum (Sera plus) for 30 min and stained with Phalloidin-TRITC (2 µg/ml, P1951, Sigma Aldrich) for 45 min at room temperature. Immunofluorescence staining for Col 1 was processed directly after phalloidin staining by overnight incubation with primary antibody (COL 1, 1:5,000 dilution with PBST, monoclonal, Sigma Aldrich). Secondary antibody Goat anti mouse, Alexa fluor 488 (Invitrogen, 1:600 diluted with PBST) was incubated for 1h. Cell nuclei were stained with DAPI (2 µg/ml, Sigma Aldrich) for 10 min. Samples were washed between every step with Tween-20 (P1379, Sigma Aldrich) 0.1% v/v in PBS and stored in PBS for microscopy.

A confocal microscope (LSM800, Carl Zeiss) was used to acquire fluorescence images (25x, 40x water-immersion objectives) to visualize Col 1 matrix and cells. Col 1 fibers within the hydrogel matrix were stained in green (λ_ex_ 488 nm), cell cytoskeleton in red (λ_ex_ 561 nm) and cell nuclei were stained with DAPI in blue (λ_ex_ 405 nm). For all samples, z-stack images were acquired for the three single channels and processed with ImageJ (National Institute of health, NIH) to generate 2D z-projection (max intensity) images. Brightness and contrast settings were adjusted to increase contrast of the single channel images. Merged images of single channels were created with the image overlay tool (merge channels) with ImageJ software (NIH). Images of Col 1 stained samples and cytoskeleton were further processed with ImageJ to quantify fiber orientation and direction of migration as described in 2.10.

#### 2.8.2 Cryo embedding

For histological staining, samples were fixed for 30 min with 4% neutral buffered formalin (Formafix AG) at room temperature and stored, washed with PBS and processed with Sucrose (150 mg/ml and 300 mg/ml, Sigma Aldrich) before being embedded in tissue freezing medium (Leica). Samples were cut with a cryostat microtome (HM 500 OM, Zeiss) to 8 µm slices and kept at -20°C until processed for histological staining with safranin O-fast green.

#### 2.8.3 Safranin O-Fast Green staining

To stain the proteoglycans in ECM within hydrogel samples and cell pellets, safranin O staining was performed after 21 days of chondrogenic differentiation. Slides were washed with water to remove the cryo-compound and stained with Weigert’s Haematoxylin (12min, room temperature, Sigma Aldrich), blued with tap water (10 min), rinsed with deionized water and stained with Fast Green (0.02% w/v, Fluka) for 6 min to visualize collagenous matrix in green/blue. After washing with acetic acid (1%, Fluka) samples were incubated with safranin O (0.1%, 15 min, Polysciences) to stain proteoglycans in red. After washing with deionized water, staining was differentiated with Ethanol (70%, Alcosuisse) and subsequently dehydrated with series of alcohols (ethanol 96%, ethanol absolute, xylene) and cover slipped (Eukitt, Sigma Aldrich). Microscopic evaluation was performed with brightfield microscope (Olympus BX63, Olympus).

### 2.9 Quantification of cell migration length and area

A sprout morphology tool in ImageJ (NIH) was used to quantify the migration length on phalloidin stained MAX projection images. For bead and sprout detection the mean threshold method was selected with sample specific adjustment of values (Bead detection: blur radius for bead detection 1.0, minimum bead radius 30-110 µm, dilate beads by factor 1.0-2.8; Sprout detection: sprout detection radius 1.0 um, minimal plexus area 5,000 µm^2^, minimal sprout area 500 µm^2^). Migration area was calculated based on resulting black and white mask from Sprout morphology tool results with ROI area measurement. The calculated area includes the area of the cell spheroids.

### 2.10 Evaluation of Col 1 fiber and cytoskeleton alignment

2D fast Fourier transform (FFT) of images acquired with SHG and confocal microscopy for Col 1 immunostaining were used for the evaluation of Col 1 fiber alignment. For both microscopic techniques, single channel images at one focus plane were used (n = 2-3 samples with 3 focus planes at the beginning, center and end of a printed filament) to investigate the homogeneity of alignment at different z-locations within the strut. For quantification of cytoskeleton orientation, MAX projection images of the red channel (561 nm) of n=4 spheroids at day 6 were processed. The Oval profile plug-in in ImageJ (NIH) was chosen for quantitative evaluation of the Col 1 fiber and cytoskeleton orientation as detailed described elsewhere [28, 29]. In brief, single images were processed with ImageJ (NIH) including unsharp mask, FFT processed, rotated 90°C right, a circular selection with radius 512 made, oval plug in for radial sum executed with number of points 360. The resulting grey values were normalized to the maximum grey values of each single image and mean values including standard deviation for both, SHG and confocal images, were shown in the bar diagram.

### 2.11 Gene expression analysis: Real-Time Quantitative Polymerase Chain Reaction (PCR)

Total RNA was isolated from the samples (THA-Col 1, THA-Col 1 MSC and MSC pellet) at day 0, 14 and 21 using TriReagent® (Molecular Research Centre Inc.) following the manufacturers’ protocol. RNA quantity was measured with NanoDrop 1000 spectrophotometer (Thermo Fischer). cDNA synthesis of 1 µg RNA was performed with Vilo Superscript (Invitrogen) according to the manufacturer’s protocol. Reverse transcription was carried out with Thermocycler (Mastercycler gradient, Eppendorf) including a pre-heating at 25°C for 10 min, followed by 42°C for 120 min, inactivation of RT for 5 min at 85°C and cooling down to 4°C.

For real time PCR, 10ul of reaction mixture containing TaqMan Universal Master Mix (Thermo Fischer), primer and probes, DEPC water and cDNA was loaded into 384 well plates. PCR was run with an initial heating to 50°C for 2 min, 95°C for 10 min, 40 cycles at 95°C for 15 s with the annealing at 60°C for 1 min.

Relative gene expression of the chondrogenic associated genes Aggrecan (*ACAN*), Col type 2 (*COL2A1*) and *SOX9*, fibrous- and hypertrophy-associated genes Col type 1 and type X (*COL1A1, COL10A1*), *RUNX2* as well as the endogenous control ubiquitin C (*UBC*) were calculated using 2^-ΔΔCq^. The *UBC* housekeeping gene has been proven for stability in the present conditions. Two different controls MSC pellets and MSC embedded in THA-Col 1 at day 0 were used for the respective samples. Details on the primers and probe sequence as well as catalogue numbers of Assay-on-Demand (Applied Biosystems) are listed in supplementary (Tab A1).

### 2.12 Quantification of Glycosaminoglycans (GAG) and DNA

To quantify GAGs and DNA within MSC pellets and cell free and cell laden THA-Col 1 hydrogels (n=3 samples/group and time point), samples were digested with Proteinase K (0.5 mg/ml, Sigma Aldrich) at 56°C. DNA content was measured in duplicates using Quant-iT™ PicoGreen (Invitrogen) assay according to the manufacturer’s instructions. Fluorescence was measured with a plate reader (Tecan infinite 200 PRO, Tecan) at 485 nm excitation and 535 nm emission including a DNA standard curve. The amount of proteoglycans was determined in duplicates by dimethylene blue dye method with absorbance measurement at 535 nm including a Chondroitin 4-sulfate sodium salt from bovine trachea (Fluka BioChemika) standard [30]. For calculation of GAGs released into media the cell free THA-Col 1 values were subtracted from the results of hMSC embedded in THA-Col 1.

### 2.13 Statistical Analysis

All samples were measured in technical duplicates of n = 2-4 biological replicates per group and time point and displayed as box plot including mean value or as mean values with standard deviation. Statistical analysis was performed with Graph Pad Prism (Prism 8, USA). Cell migration length and migration area as well as PCR data was analyzed with multiple t-test. Values of the three sample groups (THA, Col 1 and THA-Col 1) were compared at three time points (day 0, day 3 and day 8) and corrected for multiple comparisons using Holm-Sidak method. Normalization of GAG to DNA values were evaluated using two-way Anova (comparing means of each sample between day 0 and day 21 and between samples at the two time points) with Sidak post hoc test to correct for multiple comparisons. Statistical significance was assumed for p-values <0.05.

## 3 Results

### 3.1 Morphology and mechanical properties of the composite network

THA-Col 1 composites were prepared dispersing Col 1 into THA, with fibrillogenesis induced by the pH rise and THA crosslinking triggered by H_2_O_2_ addition and incubation at 37°C. The presence of Col 1 fibers in THA-Col 1 composite hydrogels was characterized by three independent techniques: SHG imaging, confocal microscopy, and turbidimetry (Fig. 2).

**Figure 2:**
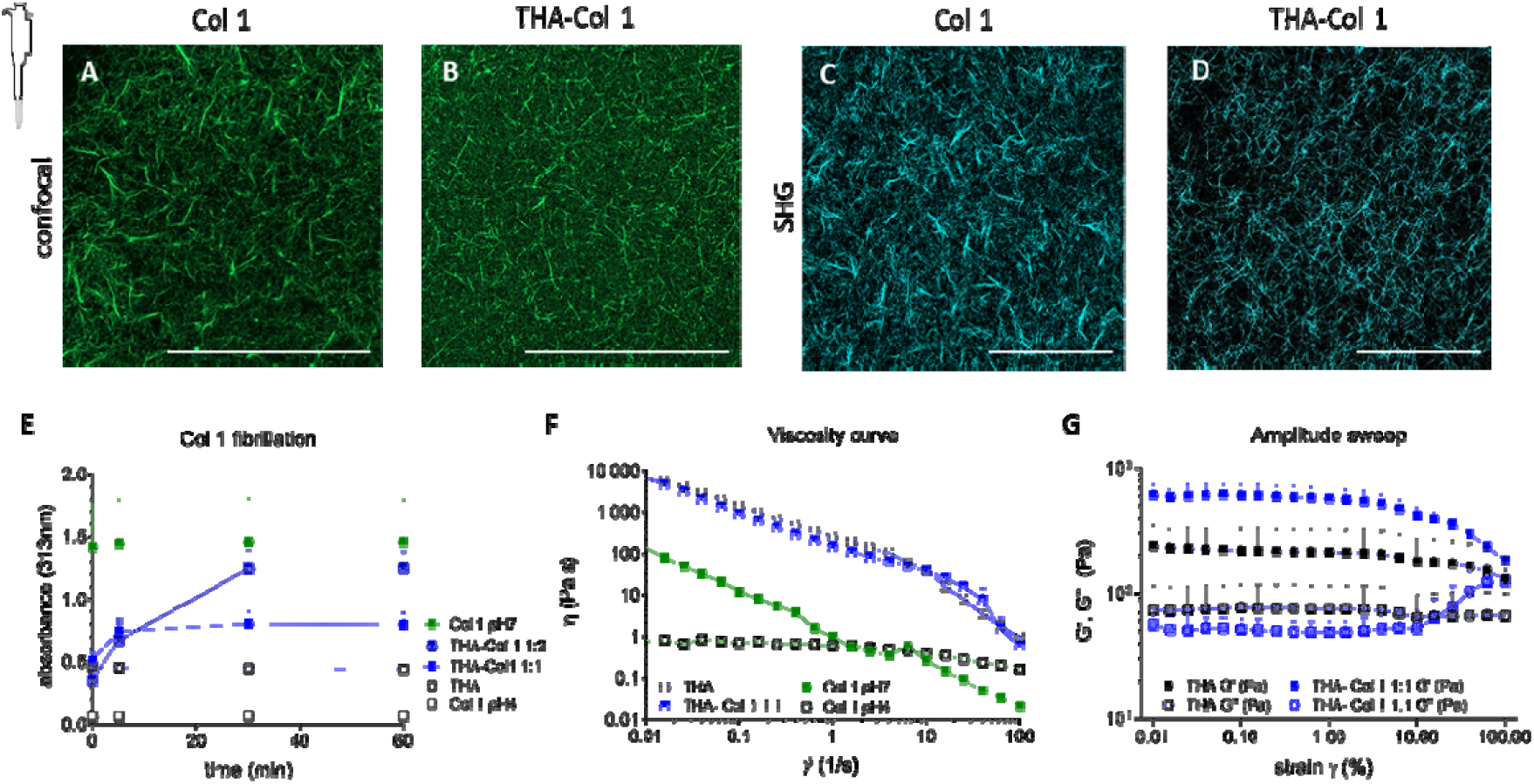
Collagen (Col 1) fibrils distribution in THA hydrogel and rheological properties. A) Visualization of Col 1 fibrils within Col 1 hydrogel and B) THA-Col 1 hydrogels using confocal imaging (immunofluorescent staining for Col 1) and C-D) SHG imaging (no labelling), respectively. Both techniques illustrate homogenous distribution of Col 1 fibrils (scale bar 100 µm). E) Turbidity measurement of THA-Col 1 demonstrated Col 1 fibril formation in composite dependent on Col 1 content. The more Col 1 is present in composite the higher the absorbance at 313 nm over time with maximum values for Col 1 only. F) Flow curve (viscosity as function of shear rate) demonstrate shear thinning behavior of THA, THA-Col 1 and neutralized Col 1 marked by decreasing viscosity with increasing shear rate. Col 1 (pH7, 5 mg/ml) showed lower viscosity compared to THA (25 mg/ml) and THA-Col 1. Acidic Col 1(pH 4, 5 mg/ml) lacks shear-thinning behavior with a constant viscosity over range of tested shear rate. G) Rheological characterization (Amplitude sweep, 1Hz) to evaluate mechanical properties of biomaterial ink by means of storage (G‘) and loss (G‘‘) modulus of THA-Col 1 and THA after enzymatic crosslinking. Storage modulus increases in composite comparted to THA.

As positive control Col 1 hydrogel was included for all measurements. The turbidity measurement allowed evaluating the fibrillogenesis kinetics (Fig. 2 E). An increase in absorbance (313 nm) over time was observed ranging from 0.4 to 0.8 for THA 1:1 Col 1 and from 0.06 to 1.3 for THA 1:2 Col 1, indicative of Col 1 fibrillation within the composite. The maximum absorbance values increased with the Col 1 content within the materials, with the highest values around 1.5 absorbance units for the neutralized Col 1 positive control, which was fibrillar from the beginning of the measurement. After 30 minutes of incubation at 37°C a plateau was reached for all groups. Pure Col 1 in acidic and in neutralized solution, and THA showed constant values in turbidity assay without change in absorbance values over time.

The fibrillation of Col 1 was further corroborated by imaging techniques. Both, SHG imaging (Fig. 2 C and D) and confocal microscopy of Col 1 immunofluorescent stained samples (Fig. 2 A and B), showed obvious presence of Col 1 fibers with a homogenous distribution within THA. Fiber morphology was visibly different, with a thinner and a denser fiber network in THA-Col 1 compared to Col 1 control. Quantification of fibril size and distribution was not possible since the fibrils cross different focal planes resulting in non-reliable values. Images acquired with confocal and SHG microscopy illustrate identical trend in fibrillar characteristics.

HRP concentration, H_2_O_2_ content and mixing ratios of THA and Col 1 were selected based on the requirements for rheological properties and extrudability of the biomaterial ink and Col 1 fibrillogenesis. The viscosity curve clearly demonstrated the shear thinning behavior of the biomaterial ink with an overall higher viscosity of 7 kPa·s for THA-Col 1 and 6 kPa·s for THA at 0.01 1/s, and monotonic decrease for increasing shear rates. At a shear rate higher than 50 1/s the viscosity of THA-Col 1 fell below the value for THA. Neutralized Col 1 (5 mg/ml) viscosity was overall lower, ranging from 0.13 k Pa·s at 0.01 1/s to 1 Pa·s at 1 1/s. Acidic Col 1 (pH=4) showed an evident reduction in shear-thinning with viscosity ranging from 0.8 Pa·s to 0.6 Pa·s. Therefore, THA-Col 1 retained the shear thinning behavior required for the biomaterial ink extrusion. An increase in storage modulus was observed in the THA-Col 1 composite hydrogel compared to THA (both samples were prepared at the same THA concentration for matching the concentrations of the enzymatic crosslinking agents, Fig. 2 G).

The amplitude sweep showed similar profile for THA and THA-Col 1, with the Col 1 presence implicating an increase in storage modulus from 220 Pa to 608 Pa at 0.1% strain, but similar decrease trend at higher strain (Fig. 2 G). This behavior is indicative that THA viscoelastic properties are not disrupted by the formation of the composite with Col 1.

### 3.2 Cell instructive properties of THA-Col 1 bioink

hMSCs spheroids were embedded into bioink made of THA-Col 1 and into its single components to determine how the composition influences cell attachment and sprouting from the spheroids. Fig. 3 A illustrates a panel of MAX projection images stained for actin filaments on day 0, day 3 and day 8. Cells migrated out of the uniformly dispersed spheroids into the biomaterials in presence of Col 1 (THA-Col 1 and Col 1), whereas no migration was observed in THA hydrogels.

**Figure 3:**
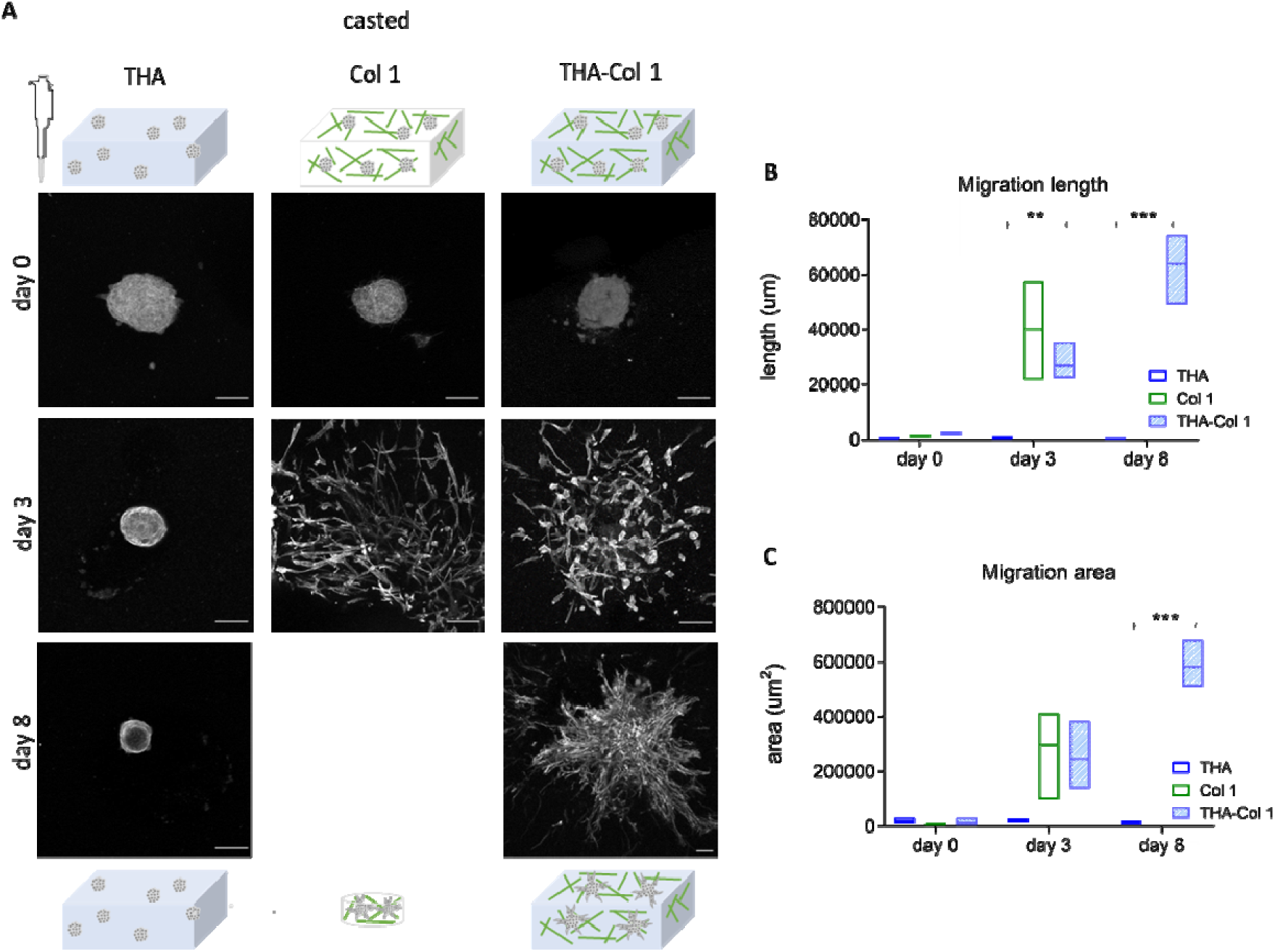
Cell instructive properties of THA-Col 1 addressing hMSC migration and cell attachment. A) Confocal images of actin filament stained hMSC spheroids embedded in THA-Col 1 (THA-Col 1, 25 mg/ml 1:1 Col 1 5 mg/ml), THA (25 mg/ml) and Col 1 (5 mg/ml) at day 0, day 3 and after 8 days culture. Migration of hMSC was seen for THA-Col 1 and Col 1 but not in THA. Col 1 hydrogel was shrinking over time and formed a small pellet of less than 2 mm in diameter (Scale bar 100 µm). B) Area of cell migration and C) migration length was higher in Col 1 compared to THA-Col 1 increasing over time. Quantification of migration area and migration length. **p<0.01, ***p<0.001.

Migration length and migration area were quantified with the ImageJ sprout Morphology Tool on MAX projections images stained with phalloidin. Mean values for both parameters confirm the highest migration in Col 1 (migration length: 40165 µm; migration area: 297 µm^2^), followed by THA-Col 1 (migration length: 26964 µm, p=0.0050 compared to THA day 3; migration area: 246 µm^2,^ p=0.0720 compared to THA day 3) on day 3. From day 3 to day 8 the migration increased further for Col 1 and THA-Col 1 (THA-Col 1 migration length: 63838 µm, p=0.0005 compared to THA day 8; migration area: 582 µm^2^, p=0.0001 compared to THA day 8). In THA, migration area and migration length of hMSC spheroids remained unvaried for the whole duration of the experiment. Due to significant shrinkage of Col 1 hydrogels the quantification of the two parameters on day 8 was not possible for for Col 1 (Fig. 3 B-C). The differences in spheroid size derived from the variation in focus plane within the 3D constructs.

### 3.3 In vitro chondrogenic differentiation

Chondrogenic differentiation potential of hMSC spheroids embedded in isotropic THA-Col 1 was compared to hMSC pellet culture as positive control, and was evaluated by qPCR, safranin-O staining and total GAG/DNA content. Gene expression analysis of hMSCs spheroid embedded in THA-Col 1 (Fig. 4 A) showed an increase in chondrogenic related markers *COL2A1* (1,000-10,000 fold), *ACAN* (10-1,000 fold) and *SOX9* (<20 fold) over time, similarly to hMSCs pellet. *COL1A1* was slightly down regulated (0.1 – 1.0-fold), while *COL10A1* showed an increase of up to 100 times for the two groups at day 14. At day 21, *COL10A1* stayed constant for hMSC embedded THA-Col 1 whereas the fold change relative to control sample decreased to <10 for hMSCs pellet. *RUNX2* upregulation was much less pronounced (< 10-fold change) compared to the extracellular matrix associated genes. The ratio *COL2A1/COL1A1* (Fig. 4 B), was higher for hMSC spheroids embedded in THA-Col 1 (day 14: 5353 ± 1699, p= 0.0242 compared to day 0, day 21: 1650 ± 409, p=0.0293 compared to day 0) compared to hMSCs pellet (day 14: 931 ± 568 p= 0.1467 compared to day 0, day 21: 300 ± 69 p= 0.0260 compared to day 0).

**Figure 4:**
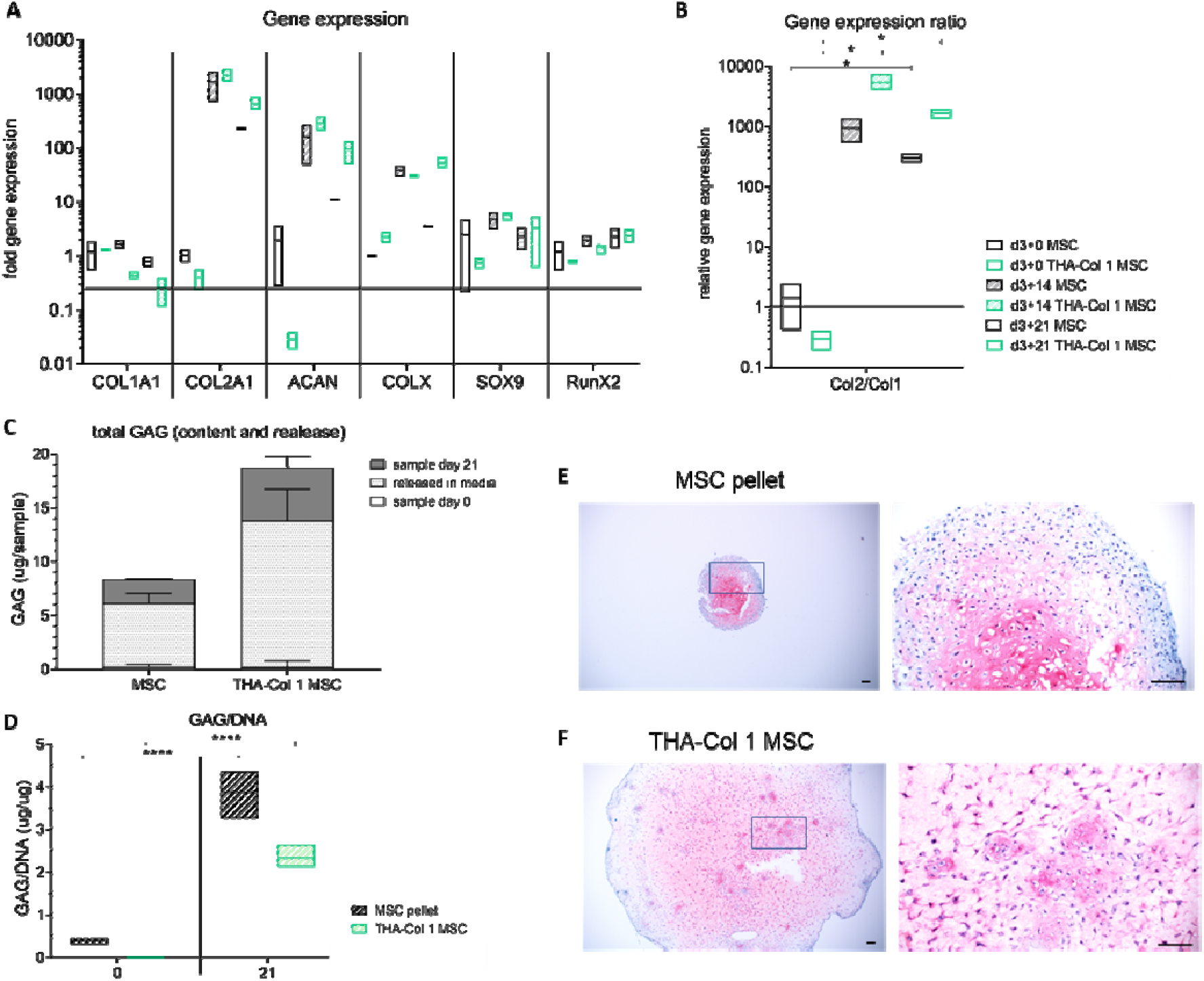
Chondrogenic differentiation potential of hMSC embedded in THA-Col 1 compared to hMSC pellet culture. A) Gene expression analysis relative to hMSC pellet or hMSC embedded in THA-Col 1 hydrogel (TC MSC) control at day 0 for Col 1 (COL 1A1), Col II (COL 2A1), aggrecan (ACAN), SOX9 and RunX2. Increase in all chondrogenic related genes with minor changes for SOX9 and RunnX2. B) Relative gene expression ratio of Col 2 to Col 1 at day 21. No statistical differences resulted for all PCR data shown beside for Col2/Col 1 ratio. *p<0.05 C) Quantification of proteoglycans (GAG/sample) including total amount of GAGs released into the media supernatant and GAGs retained in the sample at day 21. D) GAG/DNA increased over time from day 0 to day 21 for MSC pellet and THA-Col 1 MSC. **** p<0.0001. E) Safranin O staining to visualize proteoglycans in ECM after 21 days in vitro culture at two magnifications for MSC pellet and F) MSC embedded in THA-Col 1 hydrogel (THA-Col 1 MSC) (scale bar 100 µm).

At day 21, an increase in proteoglycans resulted for both groups (Fig. 4 C-D). Total amount of GAGs increased from 0.46 ± 0.38 ug/ml to 4.48 ± 0.13 ug/ml (p<0.0001) for hMSCs pellet and from 1.18 ± 0.20 ug/ml to 4.48 ± 0.69 ug/ml (p<0.0001) for hMSCs spheroids embedded in the bioink. Release of proteoglycans in the media was 5.88 ± 0.92 µg and 13.58 ± 2.93 µg, respectively. Both groups retained around 25% of the total produced proteoglycans within the sample (MSC pellet 26.8%, THA-Col 1 MSC 23.3%). Normalizing total GAG to DNA values in the samples increased for hMSCs pellet within 21 days to 3.88 ± 0.42 (p<0.0001) and for hMSCs embedded in THA-Col 1 to 2.34 ± 0.24 (p<0.0001).

The histological staining confirmed the chondrogenic differentiation. After 21 days of culture a homogenous distribution of safranin O positive staining was observed for hMSCs pellet and hMSC embedded in THA-Col 1 (Fig. 4 E-F). The hMSC spheroids were no longer visible in the point of initial seeding because of the migration throughout the whole hydrogel, resulting in a homogenous cell distribution. hMSCs morphology was comparable between the cells in the pellet and the cells embedded in the hydrogel; most cells appeared to be condensed or rounded with a low population of spindle shaped cells at day 21.

Overall, this experiment pointed out that hMSC embedded in the isotropic THA-Col 1 bioink is permissive to cell migration, GAG retention and that it retains the same chondrogenic potential as the gold standard pellet culture.

### 3.4 3DP to fabricate microscopically anisotropic scaffolds

After optimizing THA-Col 1 biomaterial ink, the Col 1 presence and fibrillar orientation were characterized. SHG imaging and confocal microscopy were used to visualize the Col 1 fibrils in the THA-Col 1 biomaterial ink after extrusion printing. The fibrillar structure of Col 1 in the composite was preserved after printing and resulted in an anisotropic material. Representative microscopic images in Fig. 5 A and C clearly demonstrate the presence of parallel aligned Col 1 fibers along the direction of the printing.

**Figure 5:**
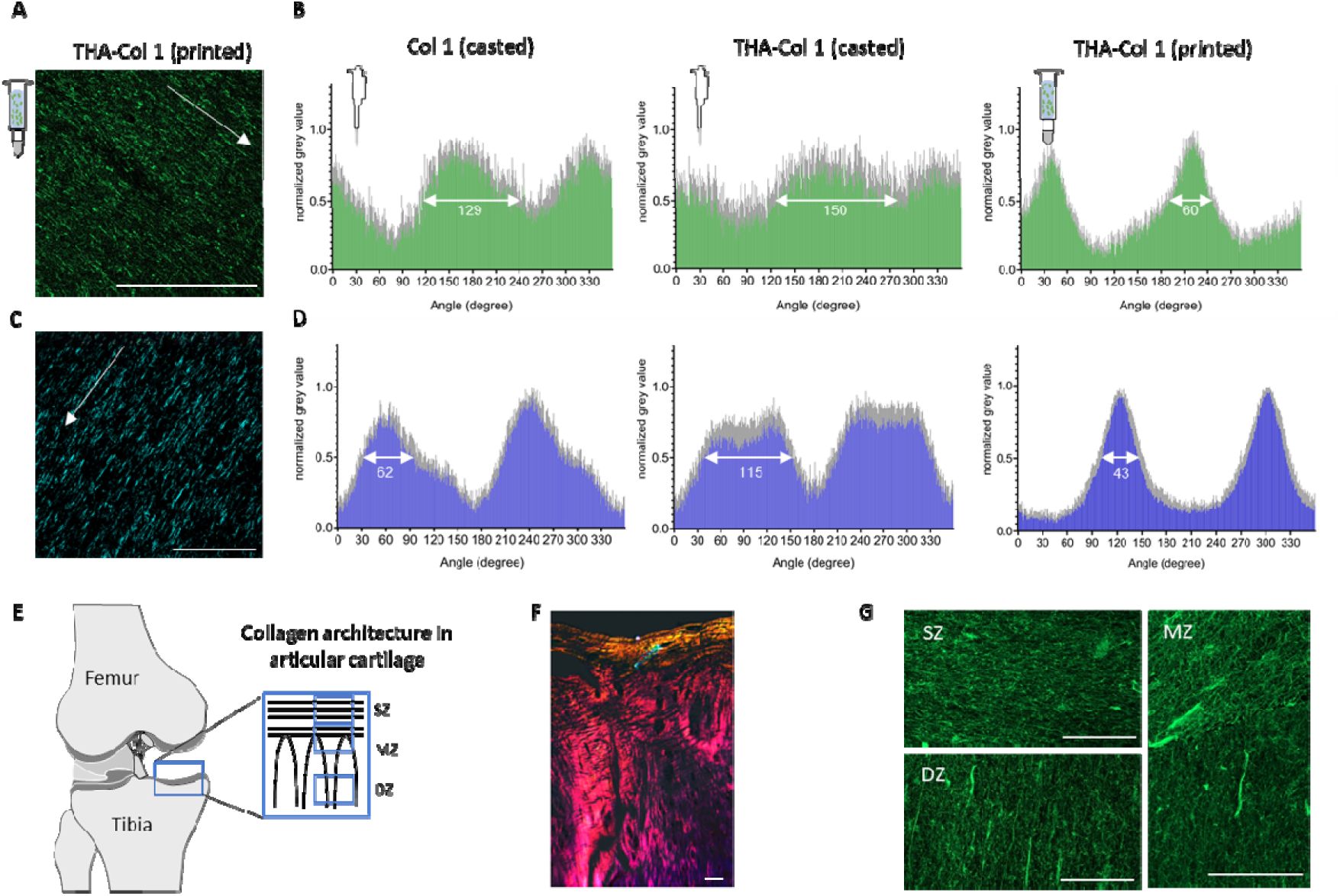
3DP as tool to align Col 1 fibers and produce microscopically anisotropic scaffolds. A) Confocal microscopy of immunofluorescent staining against Col 1 and C) SHG images demonstrate anisotropic fibers in THA-Col 1 biomaterial ink (scale bar 100 µm). Unidirectional orientation of parallel Col 1 fibers aligning along the printing direction (white arrow indicated printing direction). B, D) Bar diagrams (mean values ± standard deviation) display the quantification of fiber alignment in casted Col 1, casted THA-Col 1 (ID 1.0 mm) and printed THA-Col 1 (15G: ID 1.36 mm, 0.25” cylindrical needle) from images acquired either with confocal microscope (B, green) or SHG (D, blue). Independently of microscopical method the printed THA-Col 1 samples resulted in unidirectional orientation which was more random for casted THA-Col 1 and Col 1. White arrow in the bar diagram indicate the peak width at a normalized grey value of 0.5. E) Schematic illustration of Col 1 fiber alignment in articular cartilage. F) Polarized light image of articular cartilage illustrating horizontal fibers in superficial zone (SZ) and vertical orientation in deep zone (DZ) (scale bar 50 µm). G) Confocal images of immune-fluorescent labelled Col 1 mimicking hierarchical fiber orientation in articular cartilage with parallel horizonal fibers in the SZ, more isotropic fibers in the middle zone (MZ) and parallel vertical fibers in the DZ (scale bar 50 µm).

To further characterize this property, the ImageJ Oval plug in was used to quantify the alignment of THA-Col 1 after printing and compare to casted THA-Col 1 and casted Col 1. Casted THA-Col 1 resulted in the least anisotropic properties, followed by casted Col 1. Comparing the peak width of the preferred orientation in the printed samples at normalized grey value of 0.5 (SHG: 43 degree, confocal microscopy: 60 degree) to the corresponding casted one (SHG: 115 degree, confocal microscopy: 150 degree) and the casted Col 1 (SHG: 62 degree, confocal microscopy: 129 degree) a clear narrowing of the Col 1 fiber orientation dispersity was observed for the printed sample (Fig. 5 B and D).

The control over the parallel fiber orientation was used to produce a construct imitating the Col orientation in knee articular cartilage. Fibrillar structure in the superficial zone (SZ at the surface) was realized by printing parallel lines, whereas the Middle (MZ) and deep zone (DZ) by an arch like geometry shown in Fig. 5 E and G with an overall sample size of 1.4 × 1.6 cm. Immunofluorescent staining against Col 1 clearly shows the fibrillar alignment in the three zones with horizontal fibers in the SZ, vertical alignment in the DZ and more isotropic appearance in the MZ.

### 3.5 Effect of 3DP on cell migration in THA-Col 1 bioink

After characterization of the material microstructure and properties, we investigated the response of hMSCs on aligned anisotropic versus isotropic fibers in the THA-Col 1. The bioink was either printed with a cylindrical needle or extruded from the cartridge without needle attached and compared to casted samples. Live/dead staining of hMSC aggregates embedded in THA-Col 1 (printed versus casted) are shown in Fig. 6 A at day 1 and day 6. Cells were viable after printing and remained viable over culture time. At day 1, some dead cells (stained in red) were visible mainly in the center of the spheroids. After 6 days, less dead cells were present compared to day 1 in all three groups. The live/dead staining also showed no toxic effect of the addition of H_2_O_2_ (0.39 mM) in THA and THA-Col 1 to initiate enzymatic gelation.

**Figure 6:**
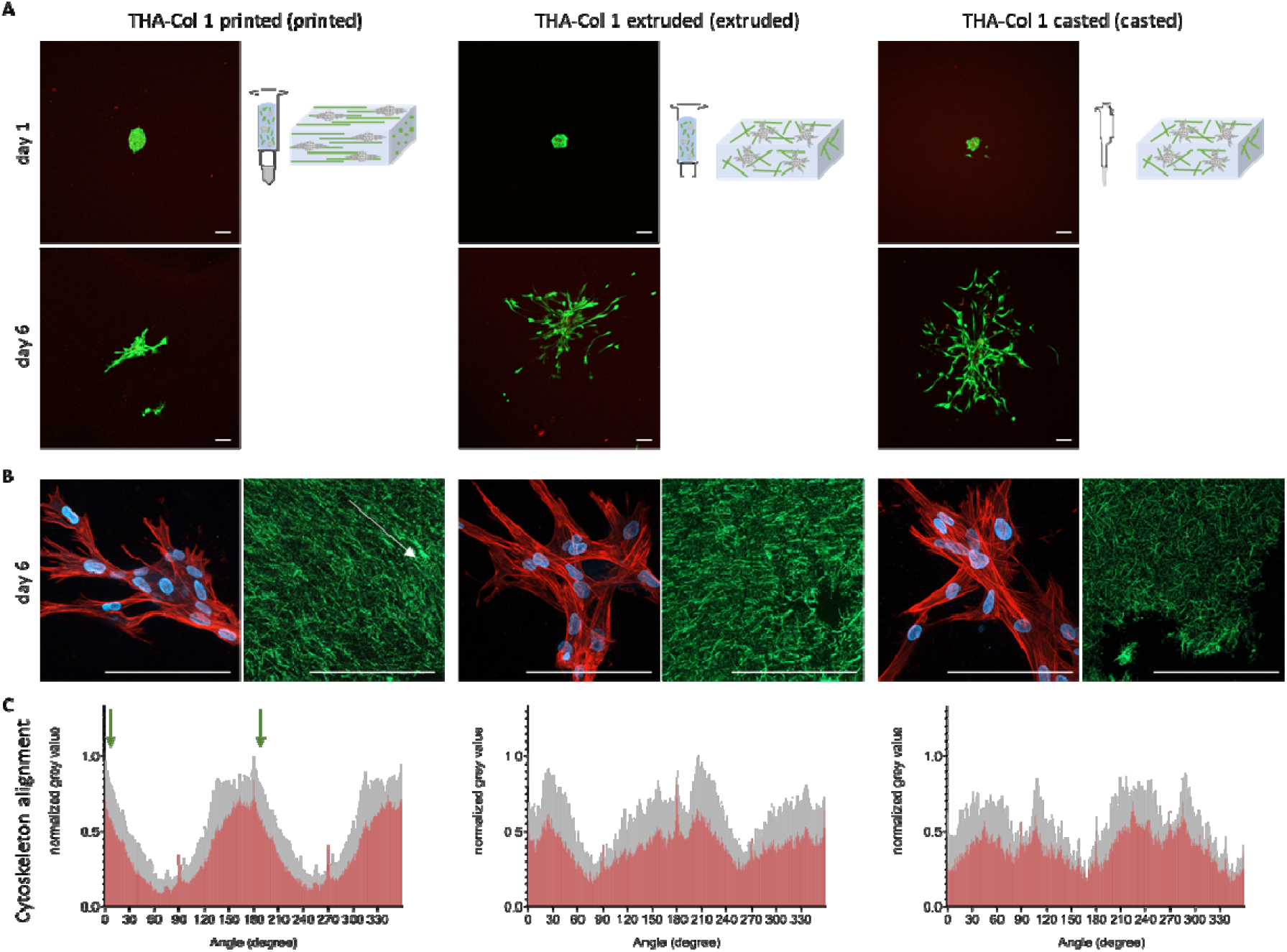
Viability (Live/dead staining) and phalloidin staining of hMSC spheroids in THA-Col 1 hydrogel. A) Representative images of live/dead staining at day 1 and day 6 showing living cells in green with few dead cells in red (Scale bar 100 µm). Bio-ink was either printed with 15G (ID 1.36 mm) cylindrical needle, extruded from 3CC cartridge (ID 2.4 mm) or pipetted with CP100 positive displacement pipette (ID 1.0 mm). B) MAX z-projection of hMSCs after 6 days of culture in THA-Col 1 demonstrating unidirectional actin filaments/cell cytoskeleton (in red) along the Col 1 (green) direction of printed samples (white arrow indicates printing direction) and cell nuclei (blue). In extruded and casted samples cells migrated random in the bioink (scale bar 100 µm). C) Orientation of cytoskeleton of hMSCs embedded in THA-Col 1 printed, extruded and casted displaying the mean values in red with standard deviation in grey. More unidirectional alignment with smaller peak width of cytoskeleton resulted for 15G printed THA-Col 1 compared to the two other groups. Green arrow in the graph of printed sample indicates the printing direction.

During *in vitro* culture, hMSC migrated from the spheroids in all three groups. Observing the cytoskeleton orientation via phalloidin staining, MSCs alignment was more pronounced in the printed groups, while hMSC showed a more isotropic cytoskeletal orientation in the extruded and casted group. The overlay image of actin filament staining (red), and cell nuclei (blue) demonstrated the unidirectional migration of the cells (Fig. 6 B).

Fig. 6 C illustrates the orientation of the cytoskeleton based on MAX projection of the red channel. The group printed with a 15G needle (left) displays two clear maxima along the orientation of the Col fibers determined by the printing direction (and the 180° from it), indicated by the green arrows, thus indicating cytoskeleton alignment along the Col fibers. For the group undergoing extrusion without needle (2.4mm diameter, center) the peaks were markedly broadened and more jagged, indicative of a more random distribution; this trend was even more apparent for the casted sample, where the distribution assumes a white noise profile.

## 4 Discussion

Extrusion-based printing has achieved significant advances in controlling construct resolution, composition and shape. However, control over the microscopic architecture has been mostly overlooked. Mechanical and biological properties of animal tissues depend not only on the chemical composition, but also on the specific spatial arrangement of structural molecules and biological factors [10]. For example, cartilage is composed of a glycosaminoglycan-based matrix containing Col 1 fibers with specific orientation, parallel to the surface on the superficial zone and perpendicular in deeper layers [31].

In the present work, we propose a technique to introduce anisotropic properties into a THA-Col 1 composite biomaterial. Having two parallel occurring crosslinking mechanisms, we overcame the difficulty in homogenous mixing Col 1 fibrils into a viscoelastic HA-based matrix by combining a previously developed bioink based on the tyramine derivative of HA [32] with acidic-solubilized Col 1, which was buffered upon mixing, thereby forming fibrils. Since both precursors, THA bioink and Col 1, are in liquid form, mixing to homogeneity was easily achieved, as visualized with microscopic techniques (Fig. 2 A-D). A distinctive advantage of the preparation method here illustrated is that THA crosslinking and Col 1 fibril formation occur simultaneously, thus avoiding phase separation [23]. Although the Col 1 fibrils in the confocal and SHG images in Figure 2 seem to differ between pure Col 1 and THA-Col 1, both casted, there is no difference in fiber diameter as shown by secondary electron and transmission electron microscopy previously [33]. The visual differences can be explained by the MAX projection of several focus plans within the 3D constructs and thus fibers in the deeper regions tend to display smaller fibrils compared to the fibrils captured in z-directions closer to the objective.

The preservation of the fibrillar microstructure of low concentrated neutral Col 1 (2.5 mg/ml final concentration in composite) within THA (12.5 mg/ml final concentration) after 3DP is a further feature allowing to produce the anisotropic microstructure.

Casted Col 1 hydrogels have been extensively used in the literature, but the fibrillar structure was not extensively investigated when combined with a second polymer. Binner *et al*. characterized a star shaped poly (ethylene glycol) hydrogel mixed with Fluorescein-isothiocyanate labelled Col 1. Besides a homogenous distribution of the Col 1, no fibrillar structure was shown and further characterized [34]. Formation of Col 1 fibers resulted to be dependent on the incubation time at 4°C prior to pH neutralization and thus inducing fibrillation when combined with Matrigel [35]. A thermally controlled printing of Col 1 (6 mg/ml) blended with Pluronic (60% w/v) resulted in Col 1 fiber formation and aggregation with a dependency of the fiber alignment with the amount of media added and thus time to dissolve the Pluronic out of the blend [21].

The shear thinning and viscosity behavior of high concentration neutralized Col 1 (20-60 mg/ml) has been characterized by different groups [20, 36]. In contrast, 3D printing of neutralized pure Col 1 at low concentrations is challenging due to the lack of shear thinning and shape retention. At the low concentration used in this study (≤ 5 mg/ml), printability was rescued by the shear thinning THA. A similar approach was presented by Duarte Capos *et al*. using agarose as shear thinning component [37]. On the other hand, printing properties of THA previously investigated in our group were preserved upon mixing with Col 1 [38].

The addition of Col 1 to THA increased the storage modulus. Thus, the fibrillar network within the viscoelastic THA can be used as a design parameter to modulate the mechanical properties. The formation of di-tyramine bonds between Col 1 and THA could be one reason for the synergistic effect [33, 39]. Other advantages of the composite are that the cell-mediated shrinking of Col 1 scaffolds is reduced; on the other side, Col 1 provides cell attachment sites to the HA, and an ECM-mimetic fibrillar matrix. Reduced shrinkage of Col 1 was also observed when this natural material was combined with silk when pulmonary fibroblasts were embedded [39].

Within the casted gels the fibrils exhibited either random orientation or local domains with some degree of orientation, which could be attributed to the low but non-zero shear forces experienced during casting from a pipette.

After 3D printing of the composite, an overall alignment along the printing axis was observed, as expected. Therefore, the low-grade enzymatic crosslinking of the THA preserved at least in part the capability of the fibrils to comply with the shear stimulus when extruded.

Shear-induced alignment of fibrillar structures has been reported by different groups [22, 40]. Kim *et al*. controlled the fiber alignment in dependence of the needle diameter during printing showing a higher degree of alignment when using 30G compared to 20G needles [40]. This intuitive behavior was not observed in the present study; rather, with thinner needles the fibrillar structure was disrupted, possibly due to the significantly higher pressure that was needed to achieve extrusion, potentially creating discontinuity or turbulence (supplementary Fig. A1). Moncal *et al*. introduced Col printing (6 mg/ml) with a thermally controlled set-up using Pluronic as sacrificial material and quantified the alignment of Col fibers along printing direction related to the amount of media and incubation time (0-48h) to induce fibrillation but with overall low anisotropy indices [21]. An agarose-Col (5 mg/ml agarose, 2 mg/ml Col 1) blend with equally distributed col fibers but no signs of alignment after printing was presented by Koepf *et al*. [41]. A different approach to align Col 1 fibers was shown by Betsch *et al*. They aligned iron nanoparticle loaded agarose-Col 1 biomaterial ink after exposure to a magnetic field. The alignment was most prevalent in Col 1 (3 mg/ml) compared to the composite (agarose 3 mg/ml, Col 1 2.5 mg/ml) only in the group with magnetic field exposure [42]. Yang *et al*. also used electrically assisted printing to mimic Col 1 orientation in meniscus by radial and circumferential aligned carbon nanotubes [43]. With the method here presented, we achieved alignment of Col 1 fibrils within a composite using exclusively natural ECM molecules avoiding using synthetic non-degradable components simply by 3D extrusion printing. Col 1 only can evolve over time becoming isotropic. This behavior is prevented within the composite, due to the presence of gelled THA stabilizing Col 1 fibrils.

The combination of the two ECM components brings biological features and directs cellular behavior. Single cell seeding resulted in enhanced adhesion, overcoming limited cell attachment reported before [38]. Similar results were observed with cell aggregates, where the presence of Col 1 increased the migration length and area. This behavior can be attributed to the presence of the integrin interaction site Arginine, Glycine and Aspartate, (RGD sequence) naturally contained in Col 1 inducing cell adhesion [44]. A direct correlation of cell adhesion and RGD density was shown also for electrospun HA functionalized with RGD peptide by Kim *et al*. [45]. The above cited agarose-Col blend has been shown to enhance smooth muscle cell spreading and attachment and thus confirming that the presence of Col 1 in a composite influences cell spreading [41].

The local microstructure of the material but also fiber parameters regulate cell response [46, 47]. In this study, hMSC migration was stimulated along the unidirectional orientation of the Col 1 fibers after 3D printing. Due to the interaction of hMSC with the Col 1 fibers, the original orientation of the parallel fiber alignment was less visible in confocal microscopy (Fig. 6 B). However, the overlapping direction of cytoskeleton orientation and fiber alignment were visualized and quantified (supplementary Fig. A2). In a comparable study, Kim *et al*. showed the unidirectional Actin filament orientation of keratinocytes embedded in corneal stroma-derived decellularized ECM after printing which were less pronounced in non-printed samples for preparation of corneal implants [40]. Yang *et al*. reported an increase in tendon associated genes of rat MSC with oriented Col 1 fiber membrane compared to the isotropic sample [48]. Whether this observation relies on the anisotropic properties or is partially induced by the shear stress during printing is not fully understood.

HA has been attributed a plethora of biological properties, including inducing cell proliferation, chondrogenic differentiation and matrix synthesis [49, 50]. The THA-Col 1 composite here proposed supported the migration and chondrogenic differentiation of hMSC spheroids, resulting in cartilage like matrix deposition throughout the whole hydrogel. Matrix deposition was uniform, with staining comparable to the standard pellet culture control (Fig. 4 E-F). Importantly, despite Col 1 presence and invasion of the whole matrix, cell morphology was also comparable to the pellet culture, showing a more condensed rather than spindle shape morphology. The biomaterial used in this study overcomes the limitation of extracellular matrix deposition only in the pericellular region which is attributed to many seminatural materials used for cartilage tissue engineering.

One possible concern regarding cartilage regeneration is the use of Col type 1 instead of type 2 for our bioink. Clinical studies of a cell free type 1 Col hydrogel (CaReS-1S^®^, Arthro Kinetics AG) implanted in focal, full layer cartilage defects have shown good clinical outcome addressing defect filling and homogenous structure of the repair tissue [51], with the Col 1 disappearing *in vivo* to leave space to the cell-deposited Col 2 [52]. In a minipig study colonization of the cell free Col 1 matrix gave outcome comparable to matrix assisted chondrocyte implantation [53, 54]. Jiang *et al*. demonstrated superior chondrogenic differentiation of MSC embedded in Col 1 compared to MSC only in a rat model [55]. On the other hand, HA has been combined with Col 1 or gelatin before with similar outcomes that chondrogenic differentiation is promoted in the composite compared to HA or Col 1 only [56-58].

One limitation of the present technique is the shear-induce alignment, limiting the degrees of spatial freedom in fibril arrangement. Another limitation is the missing quantification of the Col 1 fibers within the composite compared to Col 1 hydrogel. This was due to technical limitations in visualizing Col 1 fibrils crossing different focus planes within a 3D hydrogel and thus dealing with autofluorescence, light scattering and the opaque character of the hydrogel. For further development, the capacity of regenerating cartilage or other tissues should be tested interrogating specifically the effect of the orientation separately from the composition, comparing random oriented or non-fibrillar constructs with aligned constructs.

## 5 Conclusions

Overall, in this work, we have presented a method to obtain an THA-Col 1 composite with macroscopic homogeneity and microscopic heterogeneity mimicking the macromolecular architecture of native tissues. We achieved a uniform distribution of Col 1 fibers within an HA-based viscoelastic matrix starting from liquid precursors with simultaneous Col 1 fibrillation and HA crosslinking. The orientation of the Col 1 fibers in the 3D printed construct was controlled with the shear stress during printing and had a direct impact on cell behavior. The biomaterial ink here introduced can be extended to other fibrillar proteins to produce similar microstructural features. The possibility of printing ECM components with control over microscopic alignment brings biofabrication one step closer to capturing the complexity of biological tissues.

## Author contribution

**Andrea Schwab**: Conceptualization, Methodology, Software, Validation, Formal analysis, Investigation, Data Curation, Writing-Original Draft. **Christophe Hélary**: Conceptualization, Review and Editing. **Geoff Richards**: Review and Editing, Funding acquisition. **Mauro Alini**: Conceptualization, Review and Editing, Funding acquisition. **David Eglin**: Review and Editing, Project administration, Funding acquisition. **Matteo D’Este**: Conceptualization, Writing-Review and Editing, Supervision, Project administration, Funding acquisition.

## Declaration of competing interests

The authors declare that they have no known competing financial interests or personal relationships that could have appeared to influence the work reported in this paper.

## Data availability

The processed data required to reproduce these findings are available to download from Mendeley-Data link http://dx.doi.org/10.17632/vcxdk7s8jy.2.

## Acknowledgements

This work is part of the osteochondral defect collaborative research program supported by the AO foundation.

This project has been partially supported by “L’Agence Nationale de la Recherche” (ANR) and the Swiss National Science Foundation (SNSF): INDEED project, SNSF’s grant number 310030E_189310 and ANR’s grant number ANR-19-CE06-0028.

The authors acknowledge support of the Scientific Center for Optical and Electron Microscopy ScopeM of the Swiss Federal Institute of Technology ETHZ for acquiring SHG images.

The Graubünden Innovationsstiftung is acknowledged for its financial support.

The authors thank Dr. Christoph Sprecher for his support with confocal microscopy settings and Luca Ambrosio MD for his support setting up the turbidity measurement.

## 6 Figure captions

**Figure 7: 3D bioprinting as a tool to produce microscopic anisotropic scaffolds**. Biomaterial and bioink were prepared by mixing neutralized Col 1 (5 mg/ml) isolated from rat tails with tyramine modified HA (THA, 25 mg/ml). Neutralized Col 1 was mixed with THA for enzymatic crosslinking either cell free or containing hMSC cell spheroids. The Col 1 microstructure was investigated after 3D printing and compared to casted, isotropic samples with different microscopic techniques (Second Harmonic Generation SHG imaging and confocal microscopy). Cell instructive properties were analyzed after in vitro culture on cell migration and attachment.

**Figure 8: Collagen (Col 1) fibrils distribution in THA hydrogel and rheological properties**. A) Visualization of Col 1 fibrils within Col 1 hydrogel and B) THA-Col 1 hydrogels using confocal imaging (immunofluorescent staining for Col 1) and C-D) SHG imaging (no labelling), respectively. Both techniques illustrate homogenous distribution of Col 1 fibrils (scale bar 100 µm). E) Turbidity measurement of THA-Col 1 demonstrated Col 1 fibril formation in composite dependent on Col 1 content. The more Col 1 is present in composite the higher the absorbance at 313 nm over time with maximum values for Col 1 only. F) Flow curve (viscosity as function of shear rate) demonstrate shear thinning behavior of THA, THA-Col 1 and neutralized Col 1 marked by decreasing viscosity with increasing shear rate. Col 1 (pH7, 5 mg/ml) showed lower viscosity compared to THA (25 mg/ml) and THA-Col 1. Acidic Col 1(pH 4, 5 mg/ml) lacks shear-thinning behavior with a constant viscosity over range of tested shear rate. G) Rheological characterization (Amplitude sweep, 1Hz) to evaluate mechanical properties of biomaterial ink by means of storage (G‘) and loss (G‘‘) modulus of THA-Col 1 and THA after enzymatic crosslinking. Storage modulus increases in composite comparted to THA.

**Figure 9: Cell instructive properties of THA-Col 1 addressing hMSC migration and cell attachment**. A) Confocal images of actin filament stained hMSC spheroids embedded in THA-Col 1 (THA-Col 1, 25 mg/ml 1:1 Col 1 5 mg/ml), THA (25 mg/ml) and Col 1 (5 mg/ml) at day 0, day 3 and after 8 days culture. Migration of hMSC was seen for THA-Col 1 and Col 1 but not in THA. Col 1 hydrogel was shrinking over time and formed a small pellet of less than 2 mm in diameter (Scale bar 100 µm). B) Area of cell migration and C) migration length was higher in Col 1 compared to THA-Col 1 increasing over time. Quantification of migration area and migration length. **p<0.01, ***p<0.001.

**Figure 10: Chondrogenic differentiation potential of hMSC embedded in THA-**Col 1 **compared to hMSC pellet culture**. A) Gene expression analysis relative to hMSC pellet or hMSC embedded in THA-Col 1 hydrogel (TC MSC) control at day 0 for Col 1 (COL 1A1), Col II (COL 2A1), aggrecan (ACAN), SOX9 and RunX2. Increase in all chondrogenic related genes with minor changes for SOX9 and RunnX2. B) Relative gene expression ratio of Col 2 to Col 1 at day 21. No statistical differences resulted for all PCR data shown beside for Col2/Col 1 ratio. *p<0.05 C) Quantification of proteoglycans (GAG/sample) including total amount of GAGs released into the media supernatant and GAGs retained in the sample at day 21. D) GAG/DNA increased over time from day 0 to day 21 for MSC pellet and THA-Col 1 MSC. **** p<0.0001. E) Safranin O staining to visualize proteoglycans in ECM after 21 days in vitro culture at two magnifications for MSC pellet and F) MSC embedded in THA-Col 1 hydrogel (THA-Col 1 MSC) (scale bar 100 µm).

**Figure 11: 3DP as tool to align Col 1 fibers and produce microscopically anisotropic scaffolds**. A) Confocal microscopy of immunofluorescent staining against Col 1 and C) SHG images demonstrate anisotropic fibers in THA-Col 1 biomaterial ink (scale bar 100 µm). Unidirectional orientation of parallel Col 1 fibers aligning along the printing direction (white arrow indicated printing direction). B, D) Bar diagrams (mean values ± standard deviation) display the quantification of fiber alignment in casted Col 1, casted THA-Col 1 (ID 1.0 mm) and printed THA-Col 1 (15G: ID 1.36 mm, 0.25” cylindrical needle) from images acquired either with confocal microscope (B, green) or SHG (D, blue). Independently of microscopical method the printed THA-Col 1 samples resulted in unidirectional orientation which was more random for casted THA-Col 1 and Col 1. White arrow in the bar diagram indicate the peak width at a normalized grey value of 0.5. E) Schematic illustration of Col 1 fiber alignment in articular cartilage. F) Polarized light image of articular cartilage illustrating horizontal fibers in superficial zone (SZ) and vertical orientation in deep zone (DZ) (scale bar 50 µm). G) Confocal images of immune-fluorescent labelled Col 1 mimicking hierarchical fiber orientation in articular cartilage with parallel horizonal fibers in the SZ, more isotropic fibers in the middle zone (MZ) and parallel vertical fibers in the DZ (scale bar 50 µm).

**Figure 12: Viability (Live/dead staining) and phalloidin staining of hMSC spheroids in THA-Col 1 hydrogel**. A) Representative images of live/dead staining at day 1 and day 6 showing living cells in green with few dead cells in red (Scale bar 100 µm). Bio-ink was either printed with 15G (ID 1.36 mm) cylindrical needle, extruded from 3CC cartridge (ID 2.4 mm) or pipetted with CP100 positive displacement pipette (ID 1.0 mm). B) MAX z-projection of hMSCs after 6 days of culture in THA-Col 1 demonstrating unidirectional actin filaments/cell cytoskeleton (in red) along the Col 1 (green) direction of printed samples (white arrow indicates printing direction) and cell nuclei (blue). In extruded and casted samples cells migrated random in the bioink (scale bar 100 µm). C) Orientation of cytoskeleton of hMSCs embedded in THA-Col 1 printed, extruded and casted displaying the mean values in red with standard deviation in grey. More unidirectional alignment with smaller peak width of cytoskeleton resulted for 15G printed THA-Col 1 compared to the two other groups. Green arrow in the graph of printed sample indicates the printing direction.

## Appendix A. Supplementary information

### A.1 Supplementary Materials and Methods

#### A.1.1 3DP THA-Col 1 biomaterial ink with different needle sizes

To compare the influence of needle diameter on Col 1 fiber alignment, THA-Col 1 was extruded (3D Discovery™, RegenHU) through a 15G and 22G needle (0.25” cylindrical needles, 15G: ID 1.36 mm, 0.2 bar, feed rate 8 mm/s; 22G: ID 0.41 mm, 1.5 bar, feed rate 8 mm/s, Nordson EFD) with subsequent light crosslinking (515 nm LED, feed rate 4 mm/s). THA-Col 1 (1:1) biomaterial ink was transferred into 3CC barred (ID 2.3 mm, Nordson) for enzymatic crosslinking (30 min, 37°C), printed with the above-mentioned parameters and light crosslinked. Col 1 fibers were visualized by fluorescent microscopy after staining as described in 2.8.1. To quantify fiber orientation the Oval Plugin (imageJ, NIH) was done similar as for samples described in 2.10.

#### A.1.2 PCR: Primer details

**Table A1:**
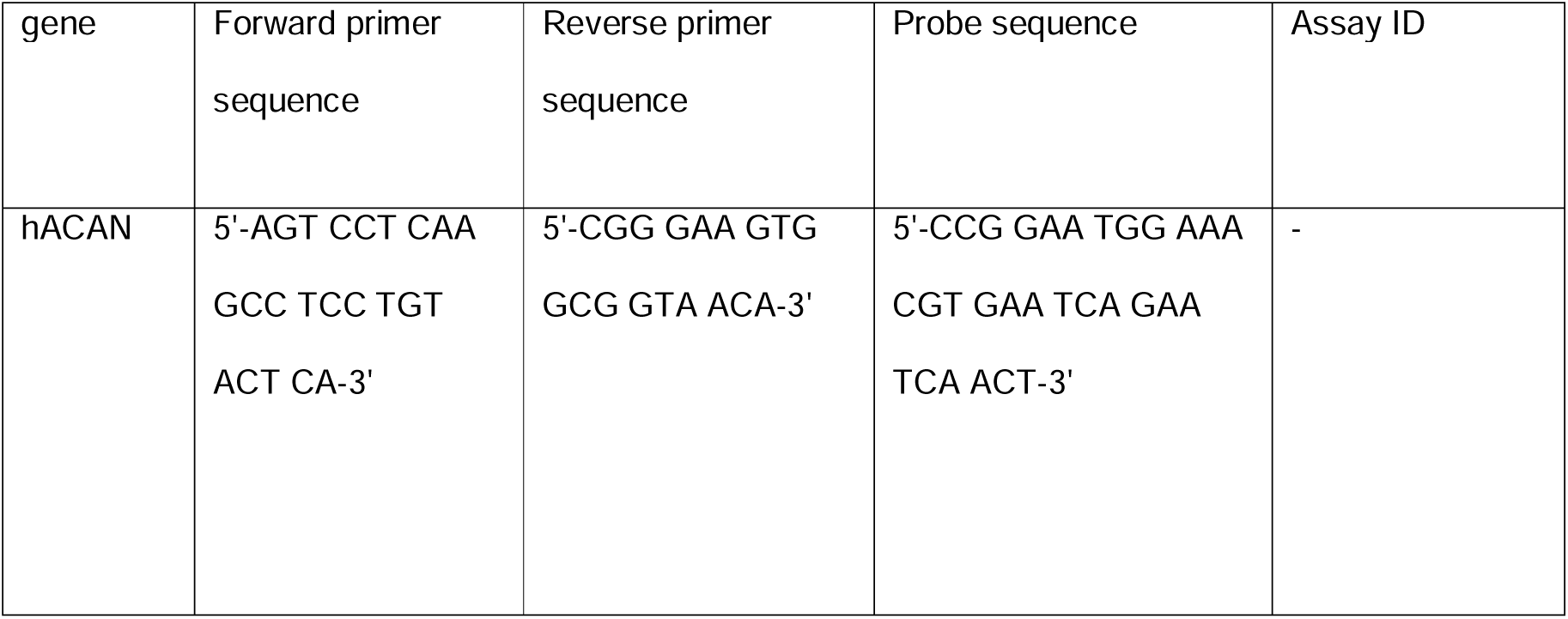
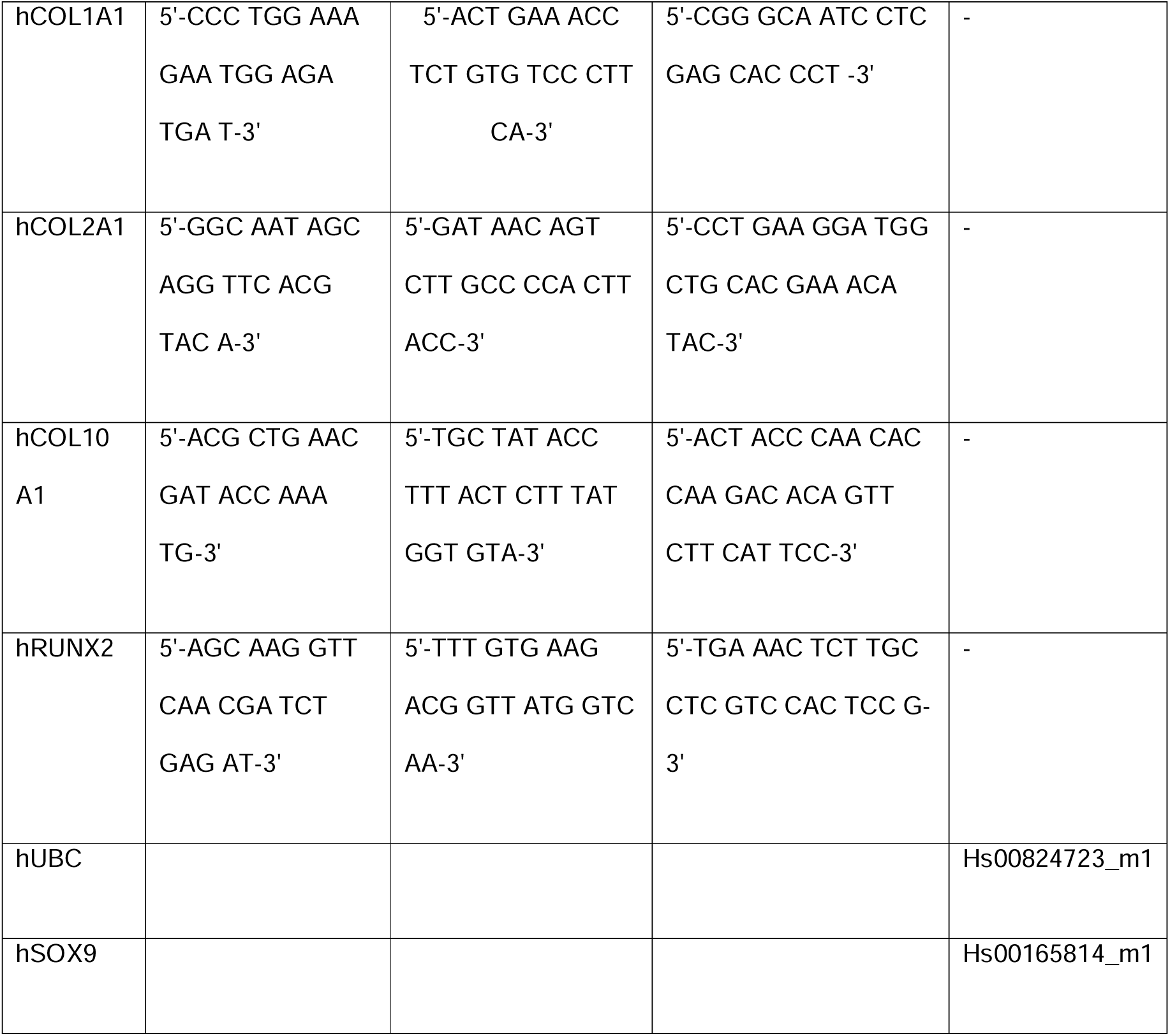
Primer and probe (Forward, reverse and probe sequence) and Assay-On-Demand (Applied biosystem Assay ID) for gene expression analysis.

### A.2 Supplementary Results

#### A.2.1 3DP THA-Col 1 biomaterial ink with different needle sizes

**Figure A1:**
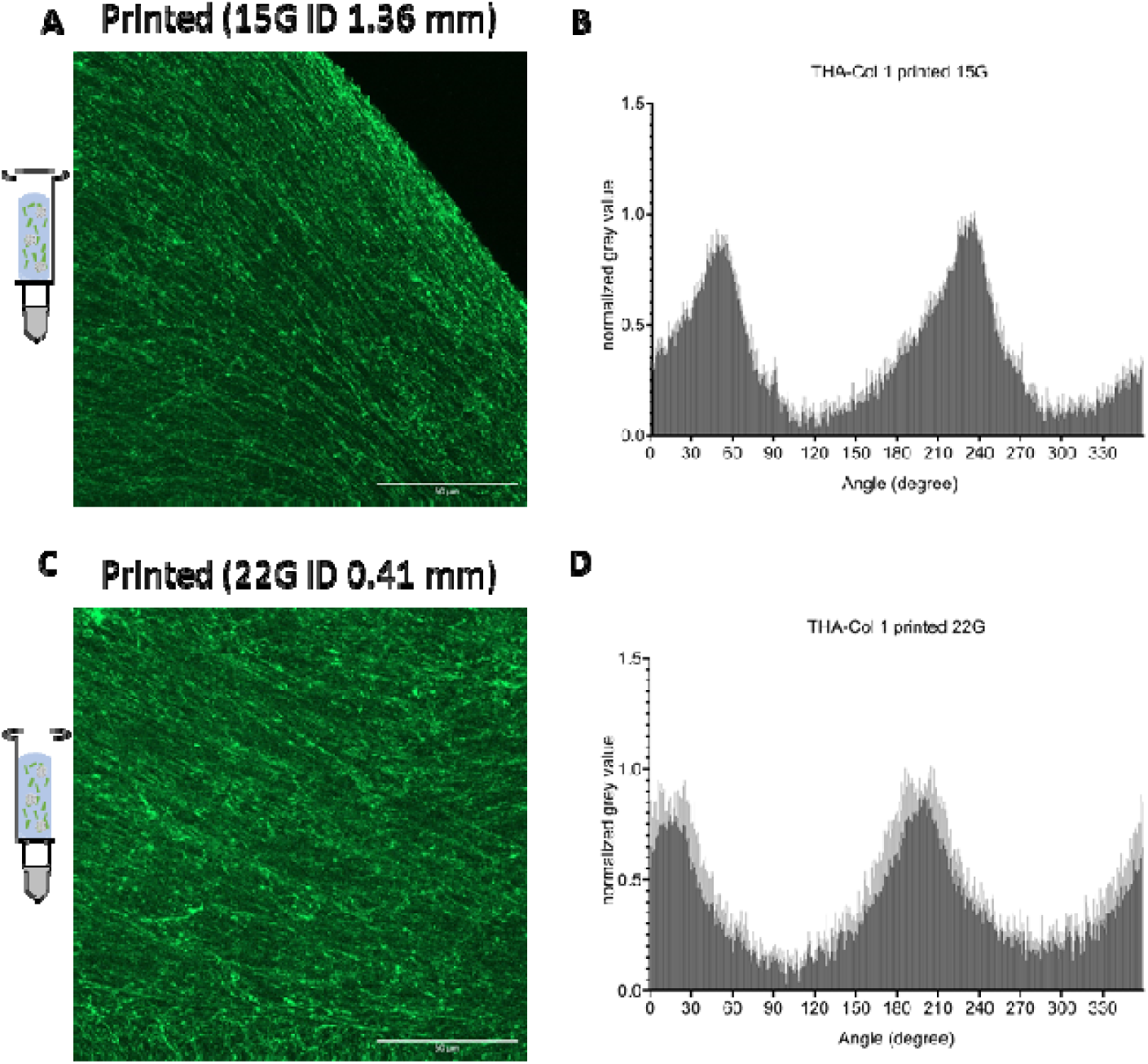
Immunofluorescent staining for collagen I of printed THA-Col 1 with 15G (left) and 22G (right) needle including printing parameters. A) Orientation of fibers did not display distinct differences. With the thinner needle diameter (22G, ID 0.41 mm) the fibers appear to become disrupted compared to the bigger needle diameter (15G, ID 1.36 mm). Scale bar 50um. B) Bar diagram displaying the quantification of fiber orientation in printed THA-Col 1 based on immune-fluorescent stained images. Peak width at normalized grey value 0.5 was smaller for 15G needle (51 degrees) compared to 22G needle (55 degrees) indicating a more heterogenous fiber orientation for the 22G needle.

#### A.2.2 3DP hMSC embedded in THA-Col 1 to analyze orientation of cell migration

Overlay of results from orientation of hMSC cytoskeleton and Col 1 fibers as described in 2.10 to illustrate the unidirectional migration of hMSC along the direction of Col 1 fibers.

**Figure A2:**
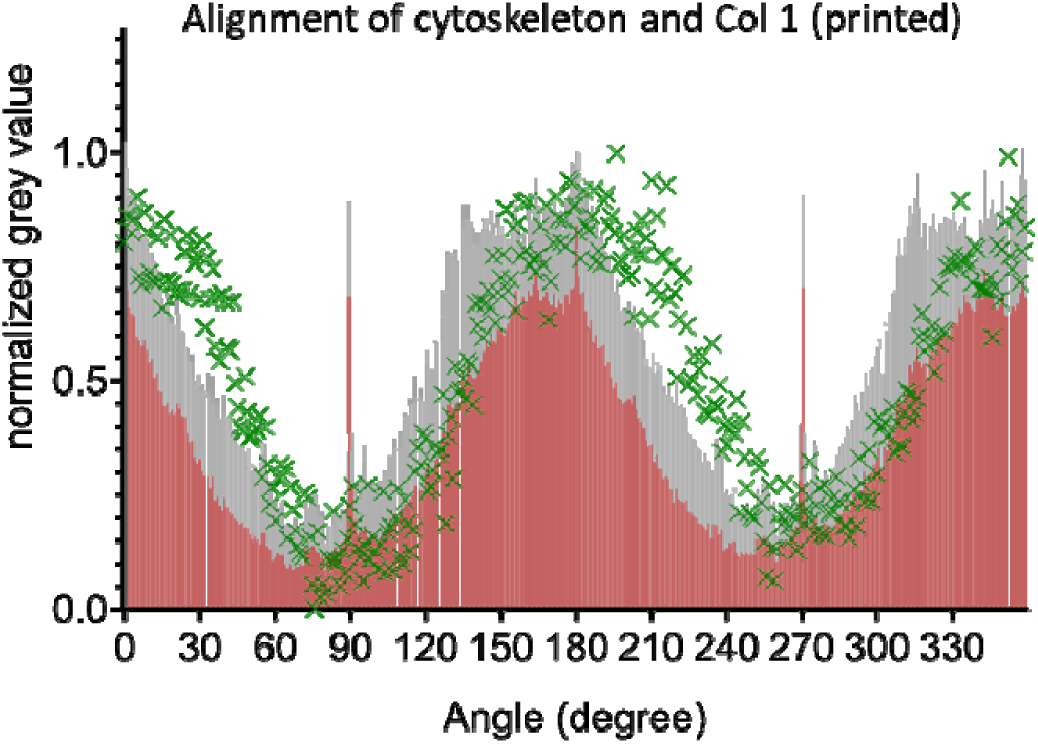
Alignment of cytoskeleton and Col 1 fibrils in printed (15G) THA-Col 1 at day 6. Overlay of mean orientation of cytoskeleton (red) with standard deviation in grey and the Col 1 orientation (green) to demonstrate the unidirectional migration of hMSCs preferably along the the direction of Col 1 fibers (grey).

## Notes

#### Summary of Updates

Graphical layout of some figures was improved, minor corrections to the text

http://dx.doi.org/10.17632/vcxdk7s8jy.2

